# Is Pisum sativum a good model species for the study of root responses to neighbours and barriers in soil? A bayesian hierarchical meta-analysis

**DOI:** 10.1101/2020.09.29.318550

**Authors:** Mariah L Mobley, Audrey S Kruse, Gordon G McNickle

## Abstract

Plant-plant competition is ubiquitous in nature. However, studying below ground behaviour of roots has always posed certain difficulties. Pea (*Pisum sativum* L.) has become a sort of model species for ecological studies about how plant roots respond to neighbouring plant roots and barriers in soil. However, published results point in several different directions. This has sometimes been interpreted as pea having sophisticated context dependent responses that can change in complex ways depending on its surroundings. To explore this further, here, we combine the result of five new experiments with published results to examine 18 unique experiments from 7 different studies for a total of 254 replicates. We used a Bayesian hierarchical meta-analysis approach to estimating the likely effect size from the available data, as well as quantify heterogeneity among different experiments, studies and cultivars. The posterior distributions show that, at the coarsest possible scale of total root production, it is unlikely that *P. sativum* root growth is influenced by either neighbours or barriers to root growth imposed by the walls of pots that vary in volume. We suggest that further work should consider repeating experiments that reported finer scale root plasticity in pea at the rhizosphere scale, and also consider alternative model species for the study of plant root responses to external cues.

## INTRODUCTION

Plant-plant competition is ubiquitous in nature, and influences individual plants, populations, communities and ecosystems (Kraft *et al.* 2015; Tilman 1982; Wilson 1988). In addition, roots must navigate through a complex soil matrix that includes neighbours, enemies, mutualists and barriers (De Deyn and Van der Putten 2005; Falik *et al.* 2005). The mechanistic details of root plasticity below ground has always been more difficult to study than above ground for reasons that are somewhat obvious (Casper and Jackson 1997; Casper *et al.* 2003; Schenk 2006). These reasons include the difficulty in getting to roots through opaque soil, the fact that most roots of most species are visually identical (Mommer *et al.* 2011; Taggart *et al.* 2011), and debate about different experimental approaches and controls (Cahill 2002; Chen *et al.* 2015; Chen *et al.* 2020; Chen *et al.* 2021; Gersani *et al.* 2001; Hess and de Kroon 2007; Laird and Aarssen 2005; McNickle 2020; Semchenko *et al.* 2010).

The common pea (*Pisum sativum* L.) has emerged as one of the most used model plants for studying root plasticity in response to varied cues in the environment by ecologists. As far as we can tell this was not a conscious choice by the research community, but rather an organic process as different groups sought to build upon previous results in the literature. Pea is an attractive model species because it has a short life history of only 50-70 days depending on cultivar, reproduces by selfing, and it is relatively small in stature. Indeed, pea has been used to study many questions about root plasticity. For example, pea has a variety of mutants that allow researchers to toggle on and off both nitrogen fixing associations and arbuscular mycorrhizal fungus association and examine the consequences of different below ground mutualisms (Guinel and Geil 2002; McNickle *et al.* 2020). Additionally, it has been shown in one study that pea roots can respond to barriers in soil, turning before contact by using exudates almost like a sonar (Falik *et al.* 2005). Pea roots have also been shown to be capable of discriminating self from non-self and exhibiting differences in root architecture when presented with self-roots or non-self-roots (Falik *et al.* 2003). In a similar study, pea was shown to preferentially proliferate into neighbour free soil volumes compared to regions of soil with more neighbouring plants (Gersani *et al.* 1998). Another study that varied nutrient dynamics in time and space concluded that pea was capable of anticipating improving conditions in the future by pre-emptively proliferating roots before the improved conditions arrived (Shemesh *et al.* 2010). At the scale of total root growth, pea has also been shown to sometimes increase total root production in the presence of both a neighbour and larger pot volume compared to alone in a smaller pot (O’Brien *et al.* 2005), and sometimes decrease root production in the same varying neighbour-volume context (Chen *et al.* 2015), or exhibit no response at all (Jacob *et al.* 2017; McNickle *et al.* 2020; Meier *et al.* 2013). Combined, these myriad results give the impression that pea has sophisticated context dependent root growth plasticity that allows complex responses to different cues.

Indeed, precisely because of these wide-ranging and interesting results in the literature we sought to further develop pea as a model for hypothesis testing about root growth plasticity and proliferation. We performed five different experiments between 2013 and 2018 where we varied different aspects of the experiment that we thought would allow us to more closely repeat some of these previous findings noted above. These used the same basic experimental design as was most common in the literature where total nutrients per plant, and nutrient concentration were controlled across plants grown alone or with neighbours (Chen *et al.* 2015; Jacob *et al.* 2017; McNickle *et al.* 2020; Meier *et al.* 2013; O’Brien *et al.* 2005). However, our experiments were not exact replications. Rather we continually fine-tuned experimental conditions in ways that we hypothesized were more similar to the conditions reported by previous authors because our prior expectation was that the published results were correct, and that we were somehow in error.

Here, we combine our five experiments with five from the literature that used the same basic experimental approach and used a hierarchical Bayesian meta-analysis approach to synthesize the results. We ask what is the average response to the barriers of pot walls imposed by changing pot volume and to the presence of neighbouring plant roots across these many experiments. We conclude that pea probably has no responses to either factor at the coarse scale of an entire root system.

## METHODS

### Studies included in the meta-analysis

We sought experiments in the published literature that grew pea plants with neighbours in pots of volume *V*, and compared it to plants grown alone in pots of volume *V*/2. This design controls total nutrients per plant, and soil nutrient concentration across the neighbour addition experiments, but has been criticised because it simultaneously manipulates pot volume and neighbours. Thus, one cannot conclude whether the barriers imposed by restricting pot volume or neighbours were the cause of any significant results (Chen *et al.* 2015; Hess and de Kroon 2007). We do not dispute this, but in the special case of no treatment effect of any kind, one can actually rule out both causes simultaneously. We identified five different studies in the literature that used this neighbour-volume experimental manipulation in pea (Chen *et al.* 2015; Jacob *et al.* 2017; McNickle *et al.* 2020; Meier *et al.* 2013; O’Brien *et al.* 2005). From these five studies we extracted the mean root and pod production within each treatment, and their standard deviations. We also recorded the cultivar used, and the pot volume used to define *V* in the neighbour-volume manipulation. In addition to the neighbour-volume manipulation, some studies imposed additional factorial treatments. These additional treatments were not replicated among studies, and so we treated these as additional treatments as separate experiments performed within study and included them as a second level of random effects in a hierarchical random effect model. These means are recorded separately resulting in multiple data points for the following studies: (i) O’Brien *et al.* (2005) crossed the neighbour addition treatment with low and high nutrient addition; (ii) Chen *et al.* (2015) included three levels of soil nutrient concentration (McNickle 2020) and; (iii) McNickle *et al.* (2020) grew plants with and without mycorrhizae. Two studies only collected root data and did not have pod data (Jacob *et al.* 2017; Meier *et al.* 2013) and one of these also did not report any estimates of variation (Jacob et al 2017).

In addition, we performed five new supplementary experiments using the same basic neighbour-volume treatment as the rest of the studies. Since these are similar to the five published results, we detail the experiments in the Supplementary information. Briefly, one of our experiments implemented the basic neighbour-volume treatment where *V* was 1L. When this did not allow us to repeat previous findings, we hypothesized that a larger rooting volume might be necessary to allow root responses. In addition, based on different approaches to allowing or preventing shoot interactions above ground by different groups (Chen *et al.* 2015; McNickle *et al.* 2020; O’Brien *et al.* 2005) we also hypothesized that shoot interactions might influence root interactions. Thus, the next four of our new experiments crossed the neighbour-volume treatment with the presence or absence of above ground shoot competition and increased the value of *V* that defined pot volumes to 6.2L. As above, we treated the presence or absence of shoot competition as separate studies nested within experiment as a hierarchical random effect and so there are actually 8 data points from these four experiments. In these four experiments, we also explored different potting media in each case (Table S1-3, Figures S1-S4).

### Meta-analysis test statistic

To compare plants in the two neighbour volume treatments, we used Hedges *g* (Hedges 1981) as our test statistic calculated according to:

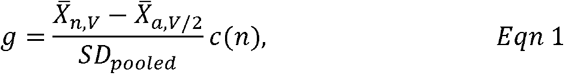

where 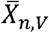 is the mean response variable in the presence of a neighbour and in a pot of volume *V* (hereafter, neighbour-full); 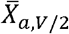 is the mean response variable when grown alone in a pot of volume *V/2* (hereafter, alone-half); *n* was the sample size of the study and *c(n)* is a correction factor for small sample size in a balanced design. The correction factor derived by Hedges (1981) for a balanced experimental design is given by:

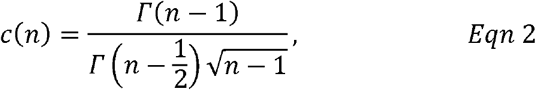

where *Γ(x)* is a gamma function of the form:

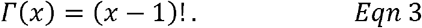

By constructing *g* with 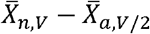 as the numerator, it will be negative in the case of reduced root growth in the neignhebour-full treatment, positive in the case of increased root growth in the neighbour-full treatment, and zero in the case of no effect of either neighbours or doubling/halving pot volume. Since all studies used a balanced design, the pooled standard deviation was calculated as:

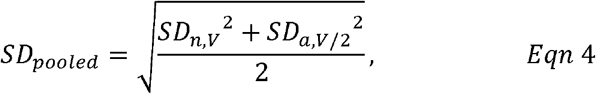

where *SD*_*n,V*_, and *SD*_*a,V/2*_ are the standard deviation associated with the means of the same subscript. We calculated *g* for individual root biomass, and also for lifetime seed yield and analysed these two tissues independently.

### Hierarchical Bayesian linear mixed effects model

A hierarchical Bayesian linear mixed effects approach was implemented with brms (Bürkner 2017; Bürkner 2018) in the R statistical environment (v 4.1.1 R-Core-Team 2021). To control for heterogeneity among studies, individual experiment was nested within study as a random effect. This allowed us to account for differences among both individual experiments (i.e. total nutrients (O’Brien *et al.* 2005), nutrient concentration (Chen *et al.* 2015; McNickle 2020), with and without mycorrhizae (McNickle *et al.* 2020), with and without shoot competition (Fig S1)), and individual research groups (E.g. soil media, fertilizer type, watering schedules, timing, and so on). Since studies also did not always use the same cultivar, and since cultivars represent a separate form of biologically interesting heterogeneity among studies in the form of genetic differences, we also included pea cultivar as another random effect. Finally, we included the pot volume used by each study as a continuous fixed effect to explicitly identify the effect of the value of *V* in the neighbour-volume response. We present both the posterior mean for each study independent of value of *V*, and the linear relationship among studies accounting for the value of *V* as a covariate. The posteriors were generated using four Markov chain Monte Carlo (MCMC) chains, 2500 burn-ins, 5,000 iterations per chain, resulting in 10,000 estimates for each posterior distribution. No thinning was used as thinning has been shown to have no detectable effects on MCMC simulation other than increased computing time (Link and Eaton 2012). To remain as unbiased as possible our priors assumed any value of *g* or standard deviation was equally likely.

## RESULTS

### Root responses

The meta-analysis included 18 unique experiments from 7 different studies for a total of 254 replicate alone-half and with neighbour-full pairs of plants. Additionally, 6 different cultivars of pea were used across the literature. The value of *V* used by studies ranged from 50mL (Meier *et al.* 2013) to 6.2 L (Fig S1).

The influence of pot volume on effect size for roots was 0.00 (95%CI: −0.12, 0.11) with an intercept of −0.03 (95% CI: −0.61, 0.58; Fig 1a). Thus, the posterior average effect size across studies was −0.03 (Table 1; Fig 2a). Unlike a frequentist approach that assigns a *p*-value to either accept or reject one hypothesis, the Bayesian framework allowed us use the posterior distributions to assign probabilities that any given outcome might be observed in a future study with peas. Here, the logical hypothesis to examine was the probability that the effect size is greater than zero (increased roots in the neighbour-full treatment) or less than zero (decreased roots in the neighbour-full treatment). Using the posterior distribution, there was a probability of 0.44 that the effect size for peas would be greater than zero, and thus a probability of 0.56 that the effect size was less than zero.

**TABLE 1:** Bayesian estimators, their error and associated 95% credible intervals (CI) from the hierarchical Bayesian mixed effects model for roots and pods in the meta-analysis. Cultivar, study and individual experiment nested within study were included as random effects. ‘Standard deviation’ is abbreviated as ‘StDev’.

| Tissue | Factor | Statistic | Estimate | Error | 95% CI |
| --- | --- | --- | --- | --- | --- |
| Roots | Mean effect size | Intercept | -0.03 | 0.28 | (-0.61, 0.58) |
|  | Pot volume | Slope | 0.00 | 0.06 | (-0.12, 0.11) |
|  | Cultivar | StDev | 0.28 | 0.26 | (0.01, 0.97) |
|  | Study | StDev | 0.28 | 0.26 | (0.01, 1.00) |
|  | Study/Experiment | StDev | 0.35 | 0.11 | (0.2, 0.6) |
| Pods | Mean effect size | Intercept | 0.05 | 0.49 | (-0.96, 1.07) |
|  | Pot volume | Slope | 0.04 | 0.07 | (-0.10, 0.19) |
|  | Cultivar | StDev | 0.51 | 0.49 | (0.02, 1.85) |
|  | Study | StDev | 0.49 | 0.52 | (0.01, 1.00) |
|  | Study/Experiment | StDev | 0.37 | 0.13 | (0.19, 0.70) |

**FIGURE 1:**
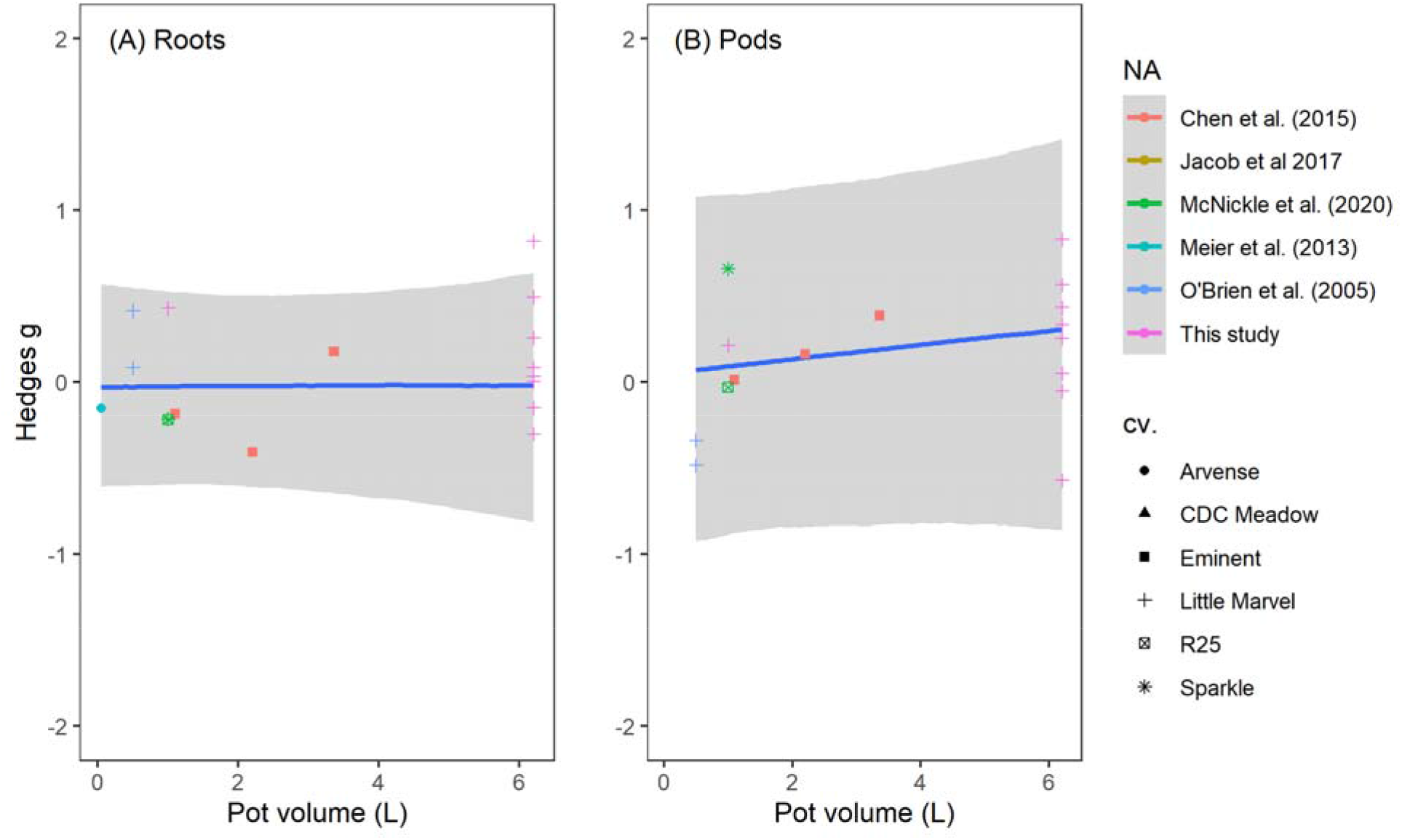
Global relationship between effect size and pot volume. Points show the observed mean effects of pot volume on log response ratio for (A) roots and (B) pods across all studies in the meta-analysis and their standard deviations. Solid lines represent the Bayesian regression line, and grey shading represents the 95% credible interval around the regression line. Cultivar, study and individual experiment within study were treated as multi-level random effects, and the heterogeneity in results introduced by these factors on the random intercept of these relationships are shown in Table 1.

**FIGURE 2:**
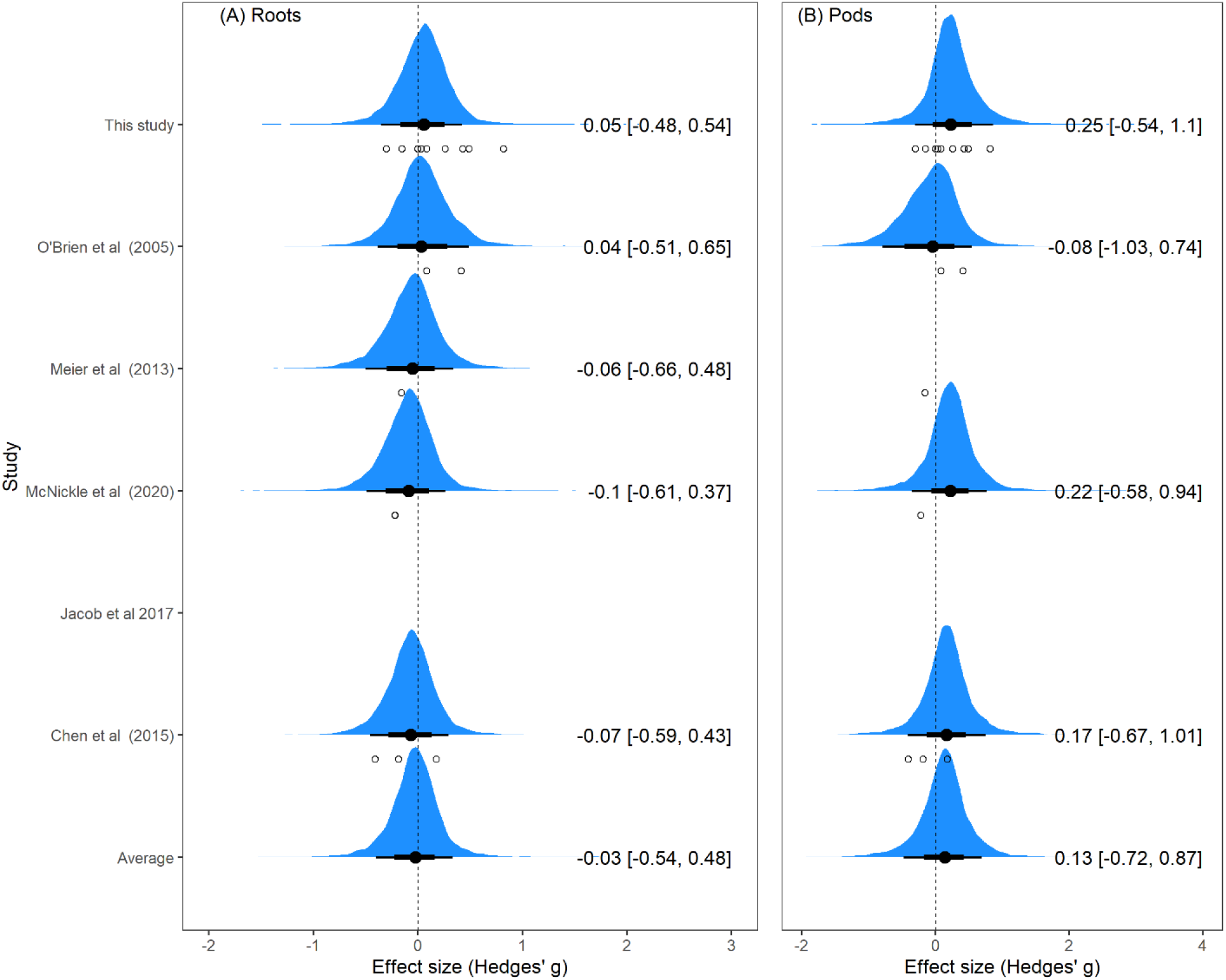
Forest plots showing the posterior distribution of effect sizes (blue) for each study and the average of all studies for (A) roots and (B) pods for each study individually (top six rows), and averaged among all studies (bottom row). Black points and lines represent posterior means with 89% (thick) and 67% (thin) credible intervals. The posterior mean and its 95% credible interval are written to the right of each distribution. White points are the observed prior effect sizes from each study. Note: since Jacob *et al.* 2017 did not report any estimate of error, and Meier *et al.* (2013) did not report error for pod mass, we cannot estimate a posterior distribution for those studies.

With any meta-analysis there is obviously differences among studies, and here quantifying that heterogeneity was major motivation for our analysis. At its core, Bayesian statistics examine how many different ways the observed data could have been sampled, and then combines these resampled outcomes into a posterior distribution. Thus, the hierarchical Bayesian mixed effects analysis is based on the assumption that each level of the random effect (experiment within study, study and cultivar) has its own effect size which emerged from resampling and are averaged to get the global result (Fig 2). Accordingly, the Bayesian approach models each of these as a standard deviation around the random intercept. Furthermore, since each random effect was modelled with its own prior distribution we can estimate this heterogeneity directly as standard deviations that also have 95% CIs (Table 1). Cultivar differences introduced an estimated standard deviation of 0.28 (95% CI: 0.01, 0.97) around the intercept, differences among studies produced an estimated standard deviation of 0.28 (95% CI: 0.01, 1.00) and differences among experiments but within a study had a standard deviation of 0.35 (95% CI: 0.2, 0.6). These should be interpreted as standard deviations around the random intercept of the model. So, since the intercept was −0.03 we can use these standard deviations to identify sources of variation that were directly associated with experimental differences, study differences and cultivar differences. The magnitude of all of these sources of error highlights that a breadth of results might be possible with a small sample size.

### Pod responses

The estimated slope of relationship between pod mass and pot volume was weakly positive at 0.04 (95% CI: −0.10, 0.19) and the intercept was 0.05 (95% CI: −0.96, 1.07; Fig 1b). Thus, pot volume increased the expected neighbour-volume effect size by 0.04 standard deviations L^−1^ of pot volume. For example, a 1L pot would have an estimated effect size of 0.09, a 2L pot would have an estimated effect size of 0.13 and so on. The average posterior effect size for most studies was also slightly positive, indicating more pod mass in the neighbour-full treatment compared to the alone-half treatment (Fig 2b). As above, we can use the posterior distributions to assign probabilities to a given outcome in pea. Here, the average posterior distribution shows that there is a probability of 0.68 that the effect size could be greater than zero, and thus a probability of 0.32 that the effect size could be less than zero in a subsequent experiment.

As above, we estimated heterogeneity in the meta-analysis directly as standard deviations around the random intercept in the model with 95% Cis (Table 1). With pods, cultivar had an estimated standard deviation of 0.51 (95% CI: 0.02, 1.85), study had a standard deviation of 0.49 (95% CI: 0.02, 1.85), and differences among experiments but within a study had a standard deviation of 0.37 (95% CI: 0.19, 0.70). Thus, there is significant uncertainty that comes along with differences in experimental design, study and cultivar used.

## DISCUSSION

Among ecologists, pea has become a sort of model species for the study of root plasticity in response to different external cues. This probably happened organically, as pea is an attractive model system, and because a number of studies had reported a variety of interesting and complex root behaviours (Chen *et al.* 2015; Chen *et al.* 2020; Falik *et al.* 2003; Falik *et al.* 2005; Gersani *et al.* 1998; Guinel and Geil 2002; Jacob *et al.* 2017; McNickle 2020; McNickle *et al.* 2020; Meier *et al.* 2013; O’Brien *et al.* 2005; Shemesh *et al.* 2010). After failing to reproduce previous findings in five of our own experiments (Supplementary information), we combined our results with those from experiments that used the same treatments to gain a more holistic picture of pea root behaviour that ultimately included 254 replicate pairs of plants grown in alone-half and neighbour full treatments. Since the literature is mostly populated by significant p-values, here we examined them in a Bayesian hierarchical meta-analysis that estimated the average posterior effect size of pea neighbour-volume effects as well as heterogeneity among studies and our priors assumed any result was equally likely. An absolute effect size of 0 < |*g*| < 0.2 is considered small, and it corresponds to just 50-58% of the treatment group being larger or smaller than the control and such a small effect would not be visually obvious. By this convention, for both roots and pods the average effect size across studies was small (−0.03 and 0.13 respectively; Fig 2). Furthermore, the 95% credible intervals around estimates was so large that – as we already knew from the literature – effectively any result might be found for root responses in any given experiment (Fig 2, Table 1). These Bayesian posterior distributions suggest that if researchers take a frequentist approach to hypothesis testing with a relatively small sample size, then whether one finds a neighbour-volume root response in pea is essentially the same as flipping a coin (Fig 2a), and only slightly better than flipping a coin that a positive pod response would occur (Fig 2b). Our original interpretation of the different results in the literature was that pea might have sophisticated context dependent responses to many cues in the rhizosphere and we sought to use this model system to explore these responses. However, based on this meta-analysis, we conclude that in general pea has neither strong neighbour root responses, nor strong responses to barriers imposed by halving pot volume when the treatments compared are two neighbour plants grown in a volume of *V*, and one plant grown in a pot of volume *V*/2.

However, in addition to the basic neighbour-full and alone-half comparison made within study, we could compare the pot volumes used among studies. The volume of the pot that peas were grown in varied from *V* = 50 mL (Meier *et al.* 2013) to *V* = 6.2 L (this study; Supplementary information) resulting in a wide range of potential barriers to root growth. However, for roots, the estimated posterior slope of this pot volume effect was 0.00 (95% CI: −0.12, 0.11; Table 1; Fig 1A). This means that one can expect the same effect size for roots whether or not plants are grown in 50mL pots or 6.2L pots. This is not the same as saying that plants did not have a growth response to pot volume (See Fig S5), only that the difference between plants in the two treatments (i.e. effect size) is zero standard deviations on average no matter the pot size that defines *V*. The influence of pot volume used in a study on the effect size for pods on the other hand was weakly positive at 0.04 (95% CI: −0.10, 0.19; Table 1; Fig 1b). This means that the effect size ranged from 0.05 in a 50mL pot to 0.30 in a 6.2L pot for pods. Thus, we conclude that the neighbour-volume manipulation does begin to have a larger effect on pod production in a positive direction, even while the treatments seem to have no effect on total root biomass. Since these experiments confound neighbour addition and pot volume it is impossible to determine which factor caused the increasing difference in pod production in neighbour-full pots relative to alone-half pots. For example, one could argue that pots of volume *V* have more nutrients and space and that this lead to the increase in pod production relative to pots of volume *V*/2 and that neighbours had nothing to do with this (e.g. Hess and de Kroon 2007). One could also argue that perhaps some kind of facilitation occurred between the two plants and that pot volume had nothing to do with the results (Callaway and Walker 1997; Thorpe *et al.* 2011). One could also argue that both neighbour and volume effects simultaneously occurred, since there is no reason to think those two ideas are mutually exclusive. These experiments cannot determine the precise cause.

However, this weakly positive slope for the effect size on pod mass (Fig 1b, Fig 2b) should also be considered in the context of the heterogeneity among experiments, studies and cultivars (Table 1). The Bayesian approach assumes that each individual experiment within study, combined with cultivar differences have their own effect size that was sampled from the global population of possible effect sizes. Therefore, these random effects produce standard deviations with their own posterior distributions that should be interpreted as standard deviations around the intercept or mean effect. Interpreting these standard deviations is easiest when considering the posterior distribution of possible effect sizes for each study (Fig 2). For example, the tails of the posterior distribution for each study include 1 and −1 for both roots and pods and thus, though the mean is centred on 0 and 0.14 respectively, there is a wide degree of uncertainty in these average estimates. For roots and pods, the uncertainty for cultivar was large which could indicate a genetic basis for the responses (Table 1). The differences among experiment performed within study indicate that other manipulations such as total nutrients (O’Brien *et al.* 2005), nutrient concentration (Chen *et al.* 2015; McNickle 2020), manipulation of mycorrhizae (McNickle *et al.* 2020), manipulation of shoot competition (Supplemental information) also influence the results, and none of this is surprising. We would direct interested readers to each study to interpret the influence of these other treatments.

Importantly, we only studied the neighbour-volume response of pea roots at the coarsest possible scale: total root biomass. Pea might have other finer scale plastic root growth in space in relation to either neighbours or barriers such as pot walls (Cabal *et al.* 2020; Falik *et al.* 2003; Falik *et al.* 2005; O’Brien *et al.* 2007). These fine-scale behaviours of individual root tips as they navigate the rhizosphere are obviously not captured by studies of total root system size. For example, Falik *et al.* (2005) found that individual root tips of pea were able to adjust growth near barriers as small as 0.8mm diameter nylon string. Such small scale behaviours would be unlikely to be detectable at the coarse scale of a total root system mass, but might still have important influences on lifetime survival and reproduction. Thus, it is still possible that finer scale root navigation was responsible for the slight differences in pod mass we observed (Fig 1b, Fig 2b). O’Brien *et al.* (2007) presented a model for finer-scale root responses to neighbours that might occur in regions of root system overlap relative to regions of soil where one plant is alone which could aide hypothesis development in future studies. Cabal *et al.* (2020) recently presented a very similar model which also makes spatially explicit predictions about intermingled root systems. If researchers want to continue to study pea responses to neighbours, we suggest that future experiments should attempt reproduce these finer scale root responses at the scale of the rhizosphere and compare them with similar approaches on other species (E.g. Belter and Cahill 2015; Downie *et al.* 2015).

Pea is an attractive model species for experimental studies of root plasticity since it is relatively small, and completes its lifecycle in 50-70 days. Our findings that pea lacks coarse scale responses does not mean that every plant on earth is expected to behave like pea. This raises the question; if one is still interested responses to neighbours and barriers in soil, then what plants might be better model species for studying root responses to neighbours?

One obvious model would be the workhorse *Arabidopsis thaliana* since there is an enormous amount of pre-existing literature, and significant genetic resources that would aide in further understanding of the neighbour-volume response (Kaul *et al.* 2000). Indeed, there is a pre-existing body of work on plant-plant competition in *Arabidopsis thaliana* (e.g. Cahill *et al.* 2005; Pantazopoulou *et al.* 2017). Another appealing option is Soybean (*Glycine max* (L.)) which also has significant genetic resources (Grant *et al.* 2009; Schmutz *et al.* 2010), and it is an economically important food crop. For such a globally important crop, even a small effect size in root or pod growth like the one detected here (Fig 1, 2) could have compounding economic implications at the scale of global agriculture and thus are worth investigating. However, as with pea there are variable results in the literature with one study finding a significant positive effect size (Gersani *et al.* 2001) and another finding an effect size of zero (Chen *et al.* 2021). These two studies used different cultivars, and so it remains to be seen if these are genetic differences, or simply the consequence of random sampling error (e.g. Fig 2). *Medicago trunata* is another common model annual plant species with significant genetic resources (Frugoli and Harris 2001) that could be considered to examine neighbour responses. Yang *et al.* (2013) found that alfalfa (*M. sativa* (L.)), a perennial congener to *M. trunata*, exhibited a root response to neighbours and so *M. trunata* might share some of these genes. However, legumes might not be the best model species for root competition studies since they form mutualisms with nitrogen fixing bacteria which might mean they have atypical root growth plasticity. Wheat (*Triticum aestivum* (L.)) might be another attractive model species with variable responses across contexts (McNickle and Evans 2018; Weiner *et al.* 2017; Zhu *et al.* 2019), significant genetic resources (Brenchley *et al.* 2012; Kidane *et al.* 2019), and economic importance as a food crop. Many other clear root responses to neighbours have been reported in herbaceous perennial plants which could be explored as new models for the neighbour response (e.g. Belter and Cahill 2015; McNickle *et al.* 2016; Padilla *et al.* 2013; Semchenko *et al.* 2010; Semchenko *et al.* 2014). However, most models include paired predictions of root growth and lifetime fitness (e.g. Gersani *et al.* 2001; McNickle and Brown 2012; McNickle and Brown 2014; O’Brien and Brown 2008; O’Brien *et al.* 2007), and so perennials might not be the best model species to test these model predictions. However, as with all scientific problems these questions will be addressed by continued experimentation over time.

### Conclusion

Common pea has been studied by ecologists in the same basic neighbour-volume manipulation across 18 unique experiments from 7 different studies for a total of 254 replicate alone-half and with neighbour-full pairs of plants. Positive, negative and neutral results have all been published and the interpretation of these results has been debated in the literature for more than a decade (Chen *et al.* 2015; Chen *et al.* 2020; Hess and de Kroon 2007; Laird and Aarssen 2005; McNickle 2020). Here, we used a hierarchical Bayesian meta-analysis to generate posterior distributions from the published literature. We find that whatever the effect of the neighbour-volume manipulation in other plant species, pea likely has no responses to a neighbour-volume manipulation at any pot volume ranging from 50mL to 6300 mL. We suggest that it might be valuable to attempt to replicate some of the finer scale results reported for pea (e.g. Falik *et al.* 2003; Falik *et al.* 2005), but for coarser scale questions it might be worth considering a new model species such as Arabidopsis, soybean, Medicago, or wheat.

## ACKNOWLEDGEMENTS

This work was supported by the USDA National Institute of Food and Agriculture Hatch project 1010722. The authors declare no conflicts of interest.

## AUTHOR CONTRIBUTIONS

GGM and MM designed the experiments. GGM performed supplemental experiment 1. MM performed supplemental experiments 2,3 and 5 and frequentist statistical analyses. AK performed supplemental experiment 4, and frequentist statistical analyses. GGM performed the Bayesian meta-analysis. MM drafted the initial manuscript, and all authors contributed to revisions.

