## Supplementary information for "Is Pisum sativum a good model species for the study of root responses to neighbours and barriers in soil? A bayesian hierarchical meta-analysis"

Supplementary methods and results for the five additional neighbour-volume experiments we performed and added to the meta-analysis. The means of root and fruit production were used to calculate effect sizes in the main text marked as “this study”.

### SUPPLEMENTARY METHODS

*Supplementary Experiment 1, V=1L*

P. sativum c.v. ‘Little Marvel’ seeds were sewn on January 22 2013 at the University of Illinois at Chicago Biology greenhouse (41°52'09.0228" N, -87°38'47.0220" W). Full sized pots were 15cm diameter, 15 cm tall circular tapered pots (V = 1 L). Plants were either grown with neighbours in pot 1L pots (hereafter neighbour-full), or alone in 0.5L pots (hereafter alone-half). In neighbour pots one of the two plants was designated ‘focal’ at the time of seeding, with the other being the ‘neighbour’. Pots were organised in a randomised block design with 10 replicate blocks. Shoot competition was controlled by pushing all pots together on the bench so that all plants experienced aboveground competition. Bamboo stakes were added next to each plant. Supplemental lighting was on a 16:8 light:dark schedule. The potting media was pure vermiculite (Coarse Vermiculite, Perlite Vermiculite Packaging Industries, Inc., North Bloomfield, Ohio, USA), and 2.5 g of slow release fertiliser (Osmocote Smart Release Plant Food Plus Outdoor & Indoor, Scotts Company LLC, Marysville, OH, USA) was added after plants germinated (Jan 28, 2013). Plants were watered every three days by measuring and pouring in 500mL of tap water and pouring it into each pot. In addition, ion exchange resin bags made from nylon mesh tubes tied on each end with string were added to the centre of each pot prior to planting. Each full sized pot received a full sized bag that contained 30mL of resin beads, while half sized pots received a half sized bag that contained 15mL of resin beads. The ion exchange resins were never analysed.

On March 23, 2013 plants were harvested by collecting fruit, shoots and roots were washed on a 2mm sieve. All tissues were dried at 60°C and weighed. In this experiment, neighbour roots could be separated during washing, and so all values represent just the focal plant. We analysed the data using generalised linear mixed effects models (GLMM) with the volume-neighbour treatment as a fixed effect, and block as a random effect. Significant differences were examined using post-hoc Tukey tests. GLMMs were performed using lme4 and lmerTest for mixed effects modeling (Bates *et al.* 2015; Kuznetsova *et al.* 2017) and lsmeans for post-hoc tests (Lenth 2016) in the R statistical environment (v. 4.1.1 R-Core-Team 2021).

*Supplementary Experiments 2-5, V=6.3L*

Peas were grown in four additional different experiments that varied in context (i.e. soil type, nutrient addition and the time of year), but included the same four neighbour treatments (Fig S1). The neighbour treatments were: 1) aboveground neighbour only; 2) below ground neighbour only; 3) no neighbour, and; 4) both above- and below ground neighbour. All experiments were performed in the same greenhouse room, and on the same bench of the Purdue University Lilly greenhouse complex, in West Lafayette, Indiana, USA (40°42'26.0"N, 86°91'88.2"W) but took place over three years. The greenhouse was set to 25°C and supplementary lighting from 600 watt high pressure sodium lights was on an 16:8 light:dark schedule for all four experiments. The same pot size was used in all experiments, with pots washed and sterilised between each experiment. To expand the range of volumes in the meta-analysis, we used large 6.2 L pots in this experiment that were 40 cm deep and 15 cm square at the top (Pot TP616, Stuewe & Sons Inc, Tanget, Oregon, USA). The planting media, and fertigation varied across the four experiments as described below. Bamboo stakes were added for each plant (i.e. two per pot).

These experiments crossed the neighbour-volume below ground treatment with and without above ground interaction. In all four experiments, above ground opaque dividers made of white corrugated plastic that were 45 cm tall and 15 cm across with 4 cm flaps to attach to the pot were constructed and placed across the middle of each pot. All pots in our experiments received dividers as a control for their effect on light reduction, and the placement of plants was adjusted such that both plants were on the same side of the divider if interaction above ground was to be permitted, or plants were on opposite sides of the divider if above ground interaction was to be blocked (Fig S1). Each treatment had two plants, which allowed us to always measure the growth of two individuals to control for potential size asymmetry (Laird and Aarssen 2005). Since the root systems of two plants were too intermingled to separate in this experiment, this two plant per treatment approach has the further advantage of always comparing the root biomass of two plants, rather than forcing us to divide the total mass of intermingled roots in half. For example, in the no-neighbour control there were still two plants, but with dividers both above- and below ground such that there was no interaction between the two plants (Fig S1; N = No neighbours), and our response variable in this treatment, as in all others, is the sum of the root system of both plants.

To minimize the effects of shading caused by the above ground dividers, and interaction among pots, replicate blocks were widely spaced on the greenhouse bench (~1m apart), and all pots were turned one-quarter turn to the east every day of the experiment. Below ground, dividers were constructed by cutting the rectangular pots in half, and nesting the two halves together (Fig S1). This created a situation where, as above, a barrier either allowed belowground interaction, or not. In all experiments, pots were arranged in a randomized block design with 15 replicates.

Each pot received two seeds five cm apart, with their location relative to the root or shoot barrier depending on treatment. Prior to sowing, the soil was saturated with tap water, and freely watered each day until germination. After germination, plants were put on strict watering regimes that differed by experiment because the different potting media had different water holding capacities and are described below (Table S1). In experiments 2, 4 and 5 fertilizer was dissolved into water and added with water on a pre-defined schedule. In experiment 3, only water was applied. The fertilizer was water soluble 24-8-16 of N-P-K solution that also contained micronutrients (Miracle-Gro All Purpose Plant Food, The Scotts Miracle-Gro Company, Marysville, Ohio, USA). The concentration and application of fertilizer varied by experiment as detailed below (Table S1).

*Experiment 2, soil:*

Experiment two was performed in early spring from February 18, 2016 to April 28, 2016. The planting media was pure potting soil (propagation mix soil, Sungro Company, Agawam, Massachusetts, USA). Each 6.2L pot was watered every other week with exactly 1L of water measured and poured into the pots (0.5L per half pot). The large pot size, and in particular the ratio of depth to surface area exposed to air, meant that soil did not completely dry between each watering. Pots were fertilized with a nutrient solution that was 0.25 g/L during weeks 3 and 6. All other experimental details were as above.

*Experiment 3, soil-gravel:*

Plants in experiment two grew extremely large and produced many fruits per individual. Thus, we hypothesized that if nutrients were highly available and not limiting to growth, plants may not experience competition because there were more nutrients than either plant could use (Casper and Jackson 1997). Thus, to reduce nutrient availability and attempt to induce competition for limited resources, the planting media in this experiment was a 1:1 mixture of potting soil and calcined clay gravel (Turface Athletics MVP, PROFILE Products LLC, Buffalo Grove, Illinois, USA). Experiment 3 was performed in early autumn from September 5, 2016 to November 14, 2016. Again, each 6.2L pot volume was watered every two weeks with exactly 1L of water. No fertilizer was applied. Unlike Experiment 2, the smaller plants in experiment 3 appeared to tip away from each other above ground because they failed to grasp the stake with their tendrils. In this experiment, bird netting (1.9 cm mesh, Bird Barricade, DeWitt Company, Sikeston, Missouri, USA) was wrapped loosely around the above ground portion of the experiment to keep plants within the vertical space above the pots. The netting was very fine and has undetectable effects on light levels (data not shown). All other experimental details were as above

*Experiment 4, gravel:*

Gersani *et al.* (1998), O'Brien *et al.* (2005) and Chen *et al.* (2015) all used nutrient free potting media in their pea experiments with pea, and applied nutrients exclusively during watering. Thus, we hypothesized that there might be something in the context of this nutrient delivery mechanism that led to their detection of neighbour responses. The planting media in experiment 4 was pure calcined clay gravel. Experiment 4 was performed in winter from December 6, 2017 to February 14, 2017. Plants were loosely tied to the stakes with stretch tie tape. The gravel did not hold water well, and so these plants were watered once each week, again by measuring exactly 1L of water per pot, however in this experiment plants were fertigated every week because the gravel did not hold water well. The fertilizer concentration was 0.5 g/L, and supplied during weeks 3, 5, 7, and 9 of growth. All other experimental details were as above.

*Experiment 5, vermiculite:*

While Gersani *et al.* (1998) and O'Brien *et al.* (2005) indeed used nutrient free potting media and nutrients supplied only in aqueous solution, they used vermiculite in both experiments, not gravel. Thus, our final hypothesis was that there was something unique about the context of growing in vermiculite compared to gravel and/or soil that might have led to their results. Thus, the planting media in experiment 5 was pure vermiculite (Coarse Vermiculite, Perlite Vermiculite Packaging Industries, Inc., North Bloomfield, Ohio, USA), which contained no nutrients. Experiment five was performed in early autumn from September 6, 2018 to November 15 2018. Plants were loosely tied to the stakes with stretch tie tape. Again, all nutrients were supplied by liquid fertilizer using the same water-fertigation schedule as in experiment 4. The other experimental details were as above.

*Harvest*

In all four experiments, after 10 weeks of growth the peas were harvested. The leaves, shoots, pods, and roots were collected separately. Roots were washed on a 2mm sieve. As expected, intermingled roots of neighbouring plants could not be separated. Tissues were dried at 60°C to constant mass, and then weighed.

*Analysis*

Since roots of interacting neighbours typically cannot not be separated, there are two approaches to dealing with this. One is to divide the total mass of plants with neighbours in half and compare this to the mass of one plant grown alone. However, this does not control for size asymmetry among the two interacting plants (Laird and Aarssen 2005). The other approach, is to pair plants growing alone and sum the biomass of two plants grown alone, thereby controlling for size asymmetry whether or not the plants actually interacted (McNickle and Brown 2014). We took this second approach. Data from each experiment was analysed using GLMM with treatment as a fixed effect, and block as a random effect using lme4 in the R statistical environment, and lsmeans to perform post-hoc tests as in experiment 1. For each experiment we examined leaf, stem, root, fruit and shoot (i.e. leaf + stem) biomass of both plants summed together as separate response variables. All biomass data was $\log(x + c)$ transformed where $c = 0.01$.

**SUPPLEMENTARY RESULTS**

*Supplementary Experiment 1, V=1L*

There was no difference in root production between plants in the alone-half or neighbours-full treatment in experiment 1 (Table S2, Fig S2A). However, plants with in the neighbours-full treatment grew significantly more shoot biomass (Table S2, Fig S2B), and significantly less fruit biomass (Table S2, Fig S2C) compared to plants grown alone.

*Supplementary Experiments 2-5, V=6.3L*

Plants in each experiment produced the same root mass (Fig S3 A-D), and pod mass (Fig S3 E-H) regardless of whether there was a neighbour above or below ground in every experiment (Table S3). Because of previously expressed concerns about pot volume effects (Chen *et al.* 2015;  Hess and de Kroon 2007), we also note that since all plants with root barriers were in pots of half the volume of all plants without root barriers, these results mean that potting volume also had no effect on plant growth in any of our experiments.

We analysed leaf and stem tissue pools separately (Fig S4). Only two significant differences arose at the $\alpha = 0.05$ significance level (Table S3). First, in experiment 2, plants that experienced root competition only, produced significantly more stem biomass than plants that experienced shoot competition only (Fig S4B). Second, in experiment 3, plants that experienced no competition produced significantly more leaf biomass than plants that experienced root competition only (Fig S4G). Though these differences were significant in the statistical sense, they were not biologically large differences. In addition, when leaf and stem mass are summed as the more commonly analysed “shoot” biomass, the significance disappeared (Table S3). Given that: (i) most root theories make no specific shoot hypotheses; (ii) the fact that the biological differences were slight (Fig 3B, G) we do not make very much of these small differences.

**TABLE S1**: Summary of methodological differences among supplementary experiments 1-5.

| | Experiment 1 | Experiment 2 | Experiment 3 | Experiment 4 | Experiment 5 |
|---|---|---|---|---|---|
| **Timing** | Winter 2013 | Spring 2016 | Autumn 2016 | Winter 2017 | Autumn 2018 |
| **Location** | Illinois 41°52'09" N, 87°38'47" W | Indiana 40°42'26" N, 86°91'88" W | Indiana 40°42'26" N, 86°91'88" W | Indiana 40°42'26" N, 86°91'88" W | Indiana 40°42'26" N, 86°91'88" W |
| **Media type** | 100% Vermiculite | 100% Soil | 50% soil 50% gravel | 100% gravel | 100% Vermiculite |
| **Volume of full pot** | 1 L | 6.3L | 6.3L | 6.3L | 6.3L |
| **Plant support** | Stakes | Stakes | Stakes and Bird netting | Stakes and ties | Stakes and ties |
| **Watering Schedule** | Every three days | Every other Week | Every other week | Weekly | Weekly |
| **Fertigation** | 2.5g slow release pellets | ¼ g/L nutrients twice | None | ½ g/L nutrients four times | ½ g/L nutrients four times |

**TABLE S2**: ANOVA table for GLMM on supplementary experiment 1. Statistical significance at the $\alpha = 0.05$ level is marked with * and bold face font. Denominator degrees of freedom (Den. df) were estimated using Satterthwaite's method.

| Tissue | Den. Df | Num. df | F | P |
|---|---|---|---|---|
| **Root** | 1 | 18 | 0.7 | 0.4035 |
| **Shoot** | 1 | 18 | 8.8 | **0.0084*** |
| **Fruit** | 1 | 18 | 35.2 | **0.0354*** |

**TABLE S3**: ANOVA tables for the GLMMs on the biomass production of different plant tissues across neighbour addition treatments in supplementary experiments 2-5. Statistical significance at the $\alpha = 0.05$ level is marked with * and bold face font. Denominator degrees of freedom (Den. df) were estimated using Satterthwaite's method.

| Tissue | Experiment | Num. df | Den. df | F | p |
|---|---|---|---|---|---|
| **Root** | **1. Soil** | 3 | 40.96 | 1.04 | 0.3852 |
| | **2. Soil-gravel** | 3 | 42.00 | 2.65 | 0.0611 |
| | **3. gravel** | 3 | 55.00 | 1.03 | 0.388 |
| | **4. Vermiculite** | 3 | 33.04 | 1.52 | 0.2276 |
| **Stem** | **1. Soil** | 3 | 41.46 | 1.57 | 0.2103 |
| | **2. Soil-gravel** | 3 | 49.00 | 3.06 | **0.0368*** |
| | **3. gravel** | 3 | 56.00 | 0.84 | 0.4761 |
| | **4. Vermiculite** | 3 | 34.64 | 1.19 | 0.3268 |
| **Leaf** | **1. Soil** | 3 | 41.01 | 0.87 | 0.4642 |
| | **2. Soil-gravel** | 3 | 49.00 | 2.43 | 0.0762 |
| | **3. gravel** | 3 | 56.00 | 3.4 | **0.0237*** |
| | **4. Vermiculite** | 3 | 34.98 | 1.09 | 0.3666 |
| **Fruit** | **1. Soil** | 3 | 41.46 | 2.02 | 0.1259 |
| | **2. Soil-gravel** | 3 | 38.6 | 0.71 | 0.5497 |
| | **3. gravel** | 3 | 42.00 | 2.79 | 0.0523 |
| | **4. Vermiculite** | 3 | 35.1 | 1.66 | 0.1939 |
| **Shoot = (Leaf + Stem)** | **1. Soil** | 3 | 41.24 | 1.25 | 0.3039 |
| | **2. Soil-gravel** | 3 | 49.00 | 2.68 | 0.0562 |
| | **3. gravel** | 3 | 56.00 | 2.47 | 0.0715 |
| | **4. Vermiculite** | 3 | 34.91 | 1.11 | 0.3561 |

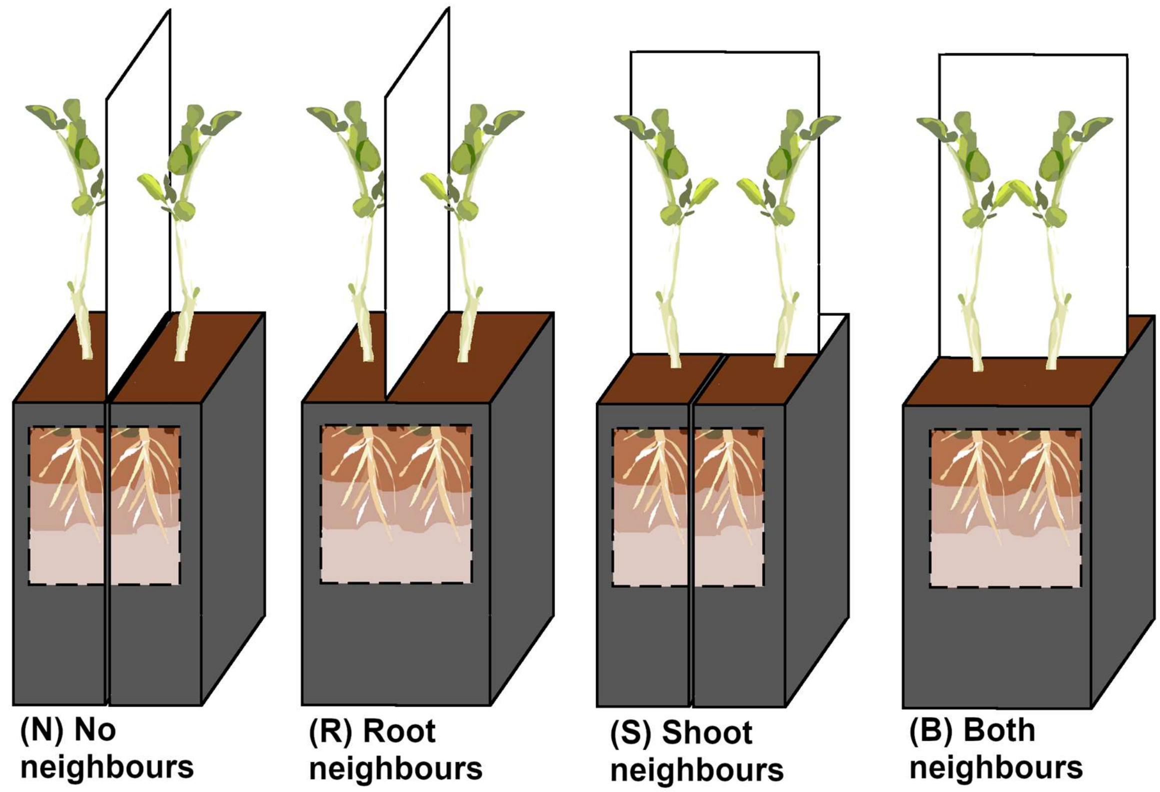


**FIGURE S1**: Schematic representation of the experimental design used to separate root and shoot competition in supplementary experiments 2-5. Dividers were placed between plants either above or belowground to create four treatments that included: (N) no interaction with neighbours; (R) only root interactions with neighbours; (S) only shoot interactions with neighbours, or; (B) both root and shoot interactions with neighbours. Image is not to scale, see methods for dimensions of pots and dividers.

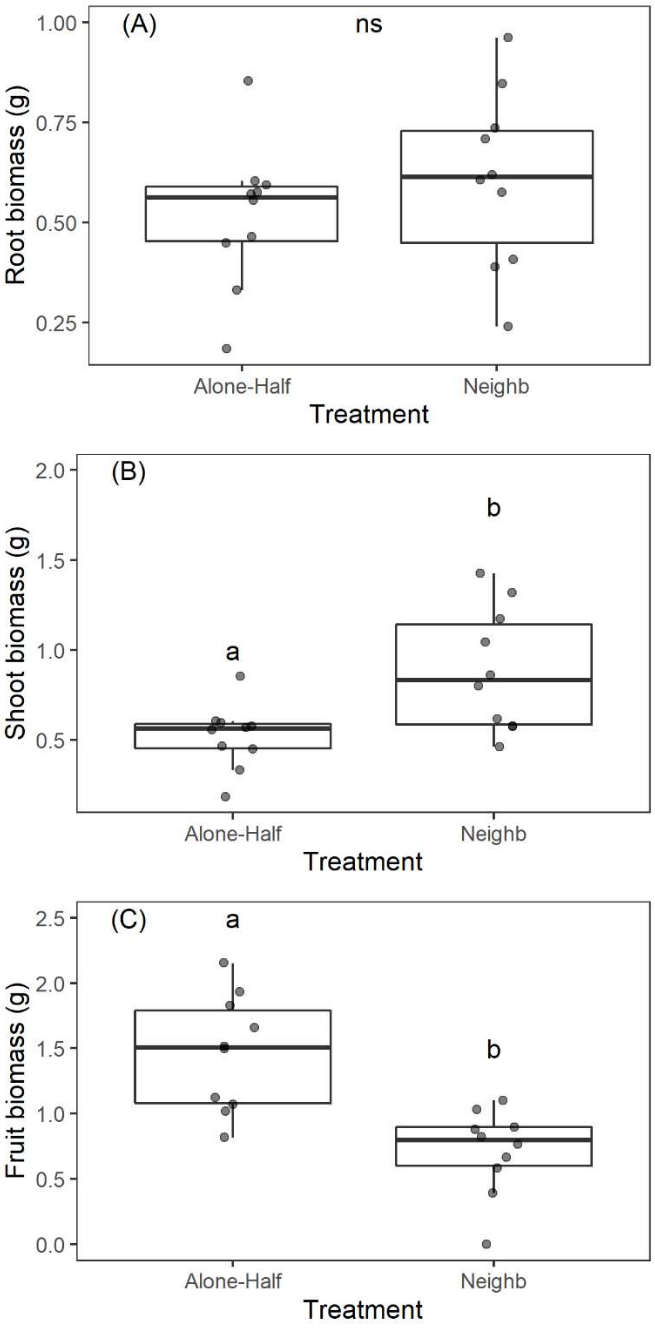


**FIGURE S2**: Biomass of plants grown alone in half sized pots (0.5L), or with neighbours in full sized pots (1L) in supplementary experiment 1. The (A) root, (B) shoot, and (C) fruit biomass of one individual plant is shown. Lower case letters indicate statistical significance, and ‘ns’ indicates lack of statistical significance at the $\alpha$=0.05 level.

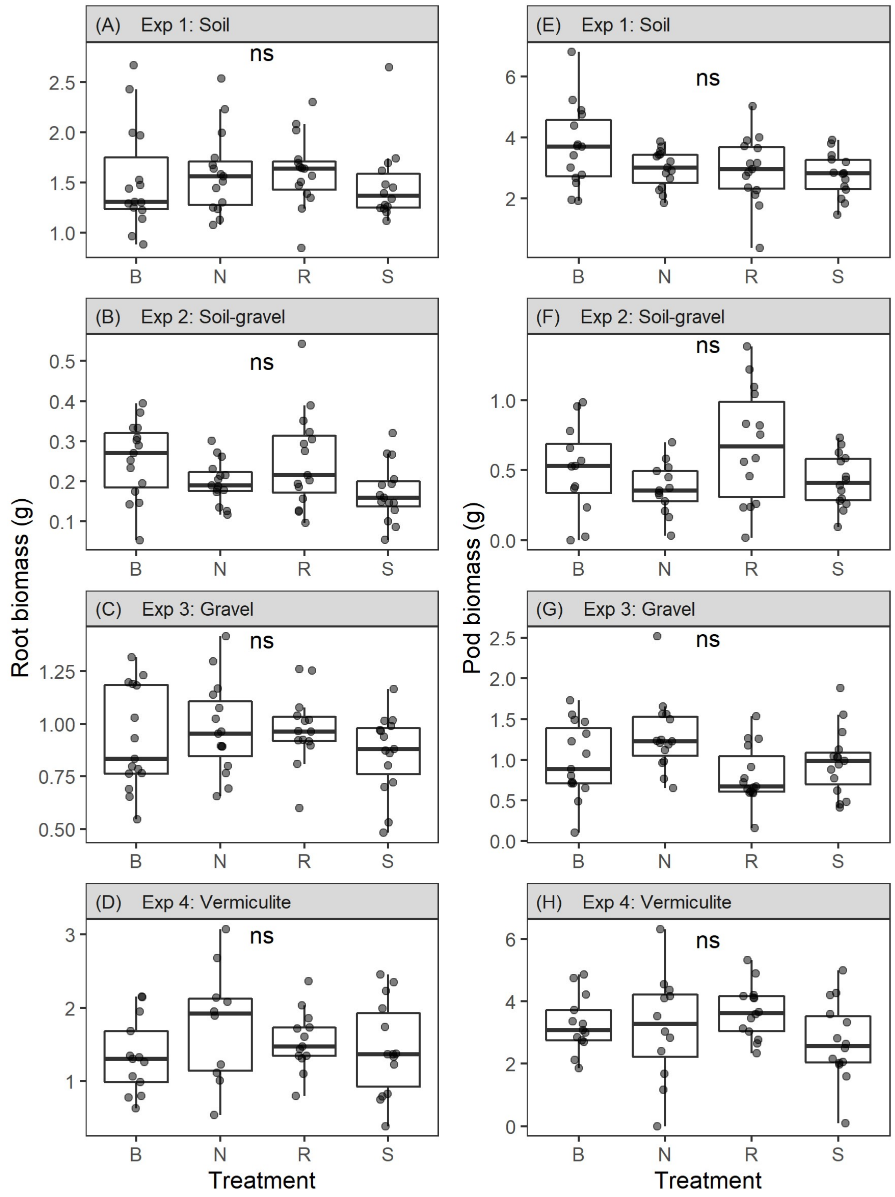


**FIGURE S3**: Results of root (A-D) and pod biomass (E-H) production across supplementary experiments 2-5. Treatment codes refer to both root and shoot interactions (B), no interactions (N), root interaction only (R), or shoot interactions only (S). The code ns indicates lack of statistical significance in GLMMs which supports the IFD model, and rejects the root under- or over-proliferation hypotheses (Table S3).

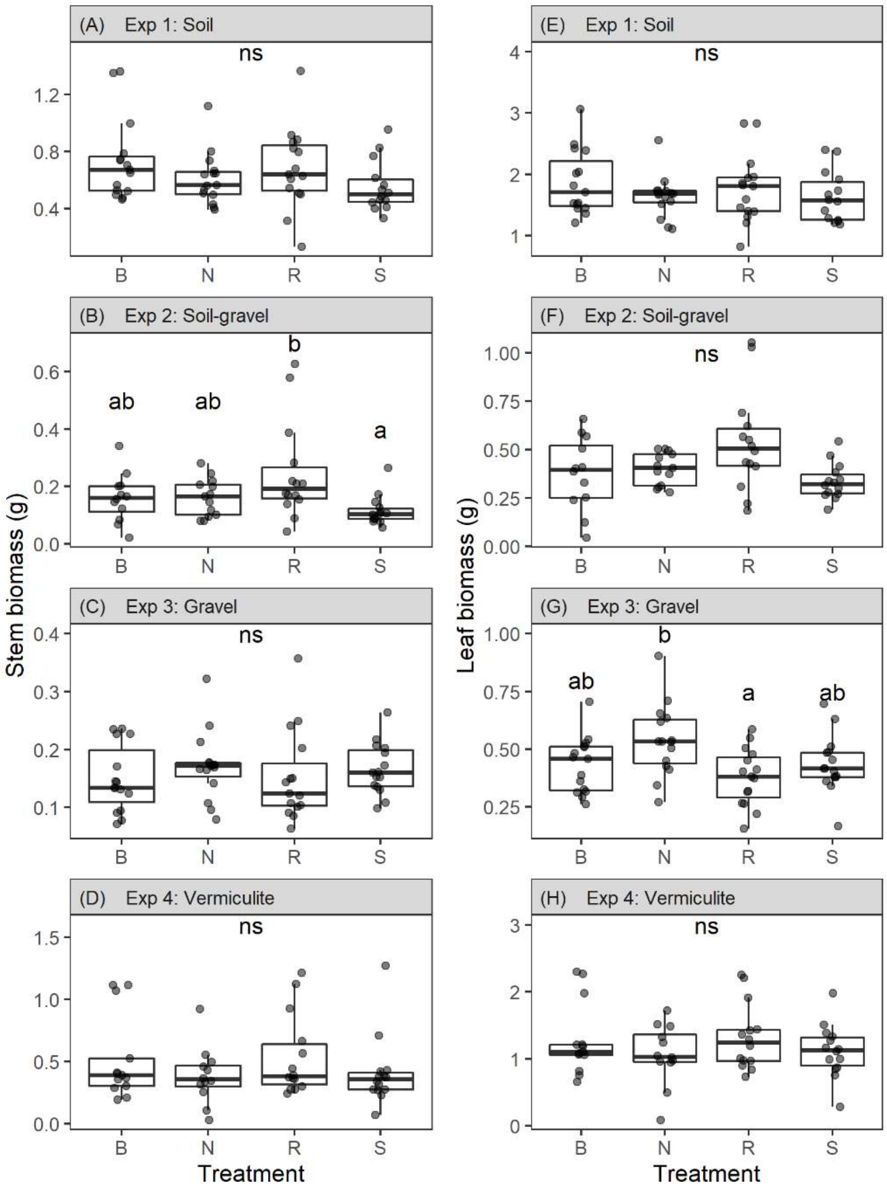


**FIGURE S4**: Results of stem and leaf production in response to neighbours either above or below ground in supplementary experiments 2-5. Most root based theories make no specific predictions about above ground growth. Treatment codes refer to both root and shoot interactions (B), no interactions (N), root interaction only (R), or shoot interactions only (S). The label *ns* indicates lack of statistical significance at the $\alpha = 0.05$ level in GLMMs, while lower case letters indicate post-hoc comparisons of differences among treatments at this $\alpha = 0.05$ level (Table S3).

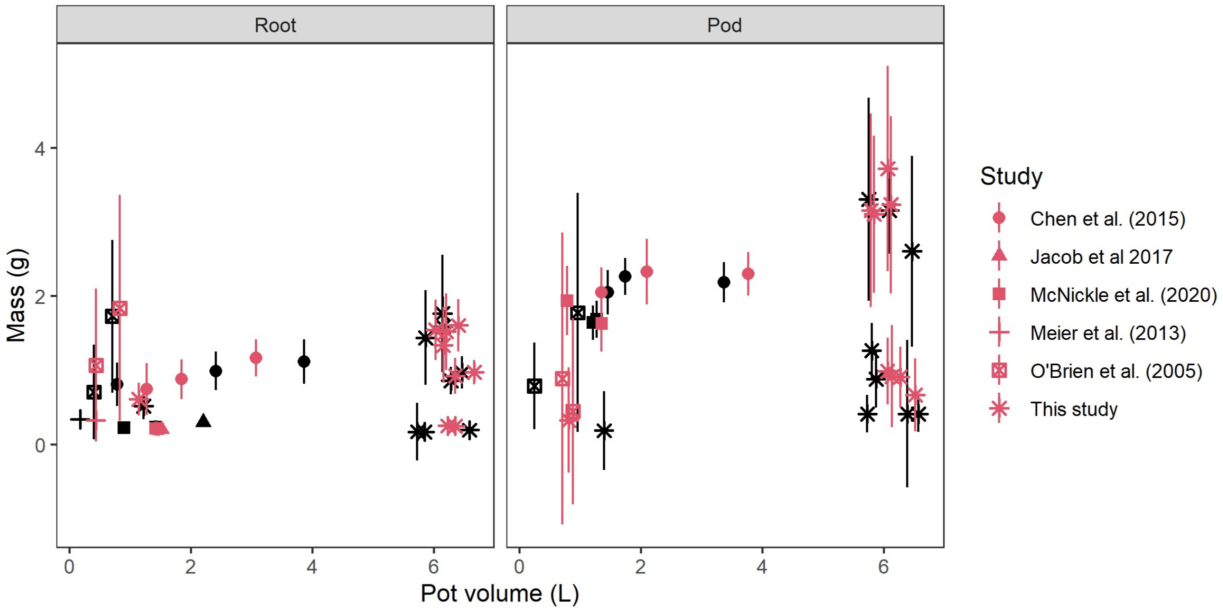


**Fig S5**: Relationship between pot volume in each study included in the meta-analysis described in the main text and raw root mass or pod mass. Plants in the alone-half treatment are shown in red, and plants in the neighbours-full treatment are shown in black. Error bars represent 1 standard deviation, and the position on the $x$-axis has a random jitter of ± 0.5 to aide in viewing studies that used the same pot volume. Linear regressions on the four relationships in this figure were not significant at the $\alpha =$ 0.05 level. Pea roots do not appear to have strong responses to pot volume, while larger volumes might lead to more pods.
